# Dissociating frontoparietal brain networks with neuroadaptive Bayesian optimization

**DOI:** 10.1101/128678

**Authors:** Romy Lorenz, Ines R. Violante, Ricardo Pio Monti, Giovanni Montana, Adam Hampshire, Robert Leech

**Affiliations:** Computational, Cognitive and Clinical Neuroscience Laboratory (C3NL), Department of Medicine, Imperial College London, London W12 0NN, UK; Department of Bioengineering, Imperial College London, London SW7 2AZ, UK; Department of Mathematics, Imperial College London, London SW7 2AZ, UK; Department of Biomedical Engineering, King’s College London, St Thomas’ Hospital, London SE1 7EH, UK

**Author notes:** These authors contributed equally to this work (Ines R. Violante, Ricardo Pio Monti).

## Abstract

Understanding the unique contributions of frontoparietal networks (FPN) in cognition is challenging because different FPNs spatially overlap and are co-activated for diverse tasks. In order to characterize these networks involves studying how they activate across many different cognitive tasks, which previously has only been possible with meta-analyses. Here, building upon meta-analyses as a starting point, we use neuroadaptive Bayesian optimization, an approach combining real-time analysis of functional neuroimaging data with machine-learning, to discover cognitive tasks that dissociate ventral and dorsal FPN activity from a large pool of tasks. We identify and subsequently refine two cognitive tasks (Deductive Reasoning and Tower of London) that are optimal for dissociating the FPNs. The identified cognitive tasks are not those predicted by meta-analysis, highlighting a different mapping between cognitive tasks and FPNs than expected. The optimization approach converged on a similar neural dissociation independently for the two different tasks, suggesting a possible common underlying functional mechanism and the need for neurally-derived cognitive taxonomies.

## Introduction

It is now well established that cognition is an emergent property of distributed networks in the brain (Bressler & Menon, 2010), and a set of frontoparietal networks (FPNs) plays a particular prominent role (Duncan, 2010, 2013). However, despite being the focus of intensive research efforts, the unique functional role of each FPN remains poorly understood (Crittenden, Mitchell, & Duncan, 2016; Hampshire & Sharp, 2015).

The most notable reason for this failure is that it is remarkably difficult to predict which cognitive tasks will isolate a FPN based on classic cognitive psychology theory (Cacioppo & Decety, 2009; Kane, Conway, Miura, & Colflesh, 2007; Poldrack & Yarkoni, 2016). The classic taxonomy of cognitive processes was developed largely blind to the functional organization of the brain; therefore, classic cognitive tasks tend to tap complex processes that involve multiple networks (Poldrack, 2006; Poldrack & Yarkoni, 2016; Price & Friston, 2005). This leads to seemingly paradoxical observations when studying the functional role of FPNs: on the one hand, FPNs are commonly co-activated during a diverse range of cognitive conditions; this is even the case when performing tasks that were originally designed to assess putatively distinct cognitive processes (Duncan & Owen, 2000; Fedorenko, Duncan, & Kanwisher, 2013; Parlatini et al., 2017). On the other hand, tasks that were originally designed to tap the same process can activate different FPNs (Hampshire, Highfield, Parkin, & Owen, 2012). This problem of overlapping functional profiles is exacerbated by the fact that FPNs also overlap spatially (Braga, Sharp, Leeson, Wise, & Leech, 2013; Erika-Florence, Leech, & Hampshire, 2014; Laird et al., 2011; Parlatini et al., 2017; Smith et al., 2009).

Resolving this many-to-many mapping (Price & Friston, 2005) problem is practically intractable with standard neuroimaging methodology because the cost and difficulty of data acquisition using functional magnetic resonance imaging (fMRI) necessitates testing only a small subset of all possible cognitive tasks. This is problematic, as studying only a fraction from the large task space can result in over-specified inferences about functional-anatomical mappings with a misleadingly narrow function being proposed as the definitive role of a network, concealing the broader role each network may play in cognition (Erika-Florence et al., 2014; Hampshire, Chamberlain, Monti, Duncan, & Owen, 2010; Hampshire & Sharp, 2015; Leech & Sharp, 2014; Shibata, Watanabe, Kawato, & Sasaki, 2016; Wager et al., 2016). In turn, this can result in inappropriately focused theories that may bias the field and potentially fuel limited generalizability and reproducibility of neuroimaging findings (Lorenz, Hampshire, & Leech, 2017; Westfall, Nichols, & Yarkoni, 2016). In the context of this problem, a more holistic understanding of the functional roles of distinct FPNs is required, and this necessitates a more comprehensive approach that examines how FPN activities vary across diverse cognitive tasks.

One such approach is to use meta-analyses, synthesizing across thousands of neuroimaging findings (Yarkoni, Poldrack, Nichols, Van Essen, & Wager, 2011; Yeo et al., 2014). While meta-analyses are powerful for tackling research questions about broad cognitive domains, they cannot extract information about fine-grained cognitive states (Yarkoni et al., 2011). In addition, they are prone to the existing biases in the literature, affected by biases in terms of which experiments were run, the reporting of results (e.g., file-drawer effect (Scargle, 2000)), selected contrasts, inconsistent labeling of brain areas and cognitive states (Poldrack, 2006; Poldrack & Yarkoni, 2016; Wager et al., 2016), as well as variable acquisition and analysis techniques (Costafreda, 2009).

To overcome these limitations in current human brain mapping methodology, we have recently proposed a radically different approach: neuroadaptive Bayesian optimization (Lorenz et al., 2016). This technique is characterized by a closed-loop search through a large task space, with fMRI data analyzed in real-time and the next task to be sampled based on the real-time results. This approach allows building upon pre-existing knowledge about the functional organization of the brain by using meta-analyses as a starting point to define a prior model of how cognitive tasks map to for example FPN activity (Lorenz et al., 2017). It then validates and refines that model in an iterative learning cycle whereby predictions are generated and experimentally tested. This produces a robustly validated model across a potentially high-dimensionality space while simultaneously identifying task conditions that produce the most optimal outcome (Brochu, Cora, & de Freitas, 2010; Shahriari, Swersky, Wang, Adams, & Freitas, 2016).

Here, we apply this approach for the first time for discovering cognitive tasks that maximally dissociate dorsal (dFPN) from ventral FPNs (vFPN); both of which are core parts of the multiple demand cortex and known to co-activate across diverse cognitive conditions (Duncan & Owen, 2000).

We achieved this in three-stages (Fig. 1) including: (1) previous meta-analysis to identify tasks that recruit these networks (Yeo et al., 2014), (2) real-time mapping of the task-network space to select optimal tasks that dissociate networks and (3) real-time optimization of the selected tasks in order to maximize the dissociations.

**Figure 1:**
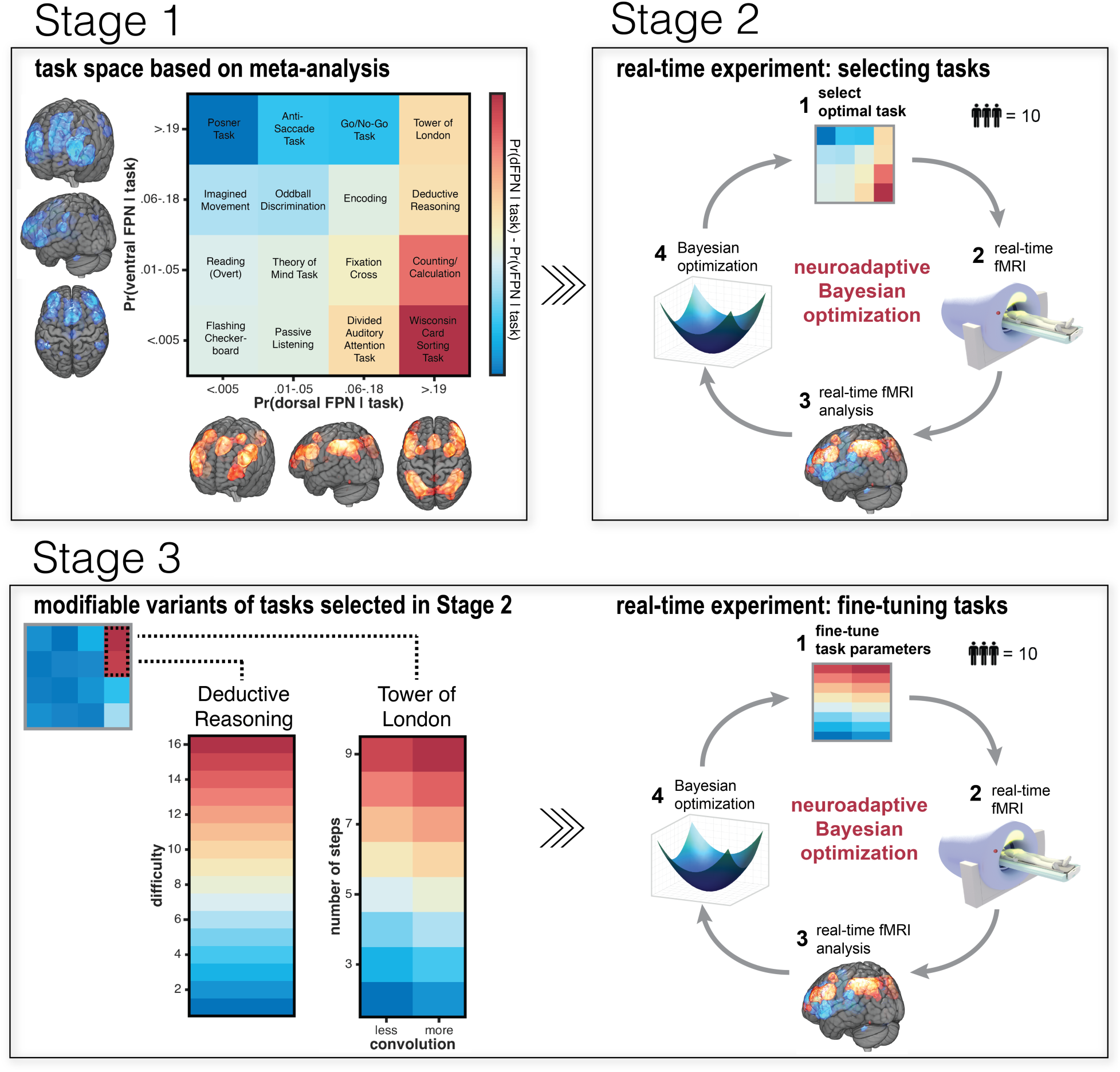
Overview of methodology. In **Stage 1,** a 2D-task space was designed with each dimension corresponding to the weighting of 16 tasks on the dFPN and vFPN, derived from a previous meta-analysis (Yeo et al., 2014). Color-coding indicates hypothesized difference for dFPN>vFPN. In **Stage 2** this task space was searched by the optimization algorithm to find optimal tasks that dissociate networks (i.e., dFPN > vFPN). In **Stage 3,** task parameters of optimal tasks from Stage 2 were fine-tuned. For Deductive Reasoning task, the optimization algorithm searched across 1D-space with 16×1 difficulty levels. For Tower of London task, the optimization algorithm searched across 2D-space with 8 (number of steps) × 2 (convolution) parameters.

## Results

In Stage 1, a 2D-task space was designed based on a previous meta-analysis (Yeo et al., 2014), with each dimension corresponding to the weighting of 16 different cognitive tasks on the dFPN and vFPN (Fig. 1). For Stage 2 and Stage 3, we employed a closed-loop neuroadaptive Bayesian optimization approach as described in Lorenz et al. (Lorenz et al., 2016). Bayesian optimization is characterized by a *surrogate* model that captures the algorithm’s beliefs about the unknown relationship between the task space and the subject’s brain response. Commonly, a Gaussian process (GP)-based model is used due to its versatility and flexibility (Rasmussen & Williams, 2006). The approach is iterative; subjects are presented with an experimental condition (e.g., one of the 16 different tasks for Stage 2 or different task parameters in Stage 3) and their brain response is measured in real-time. This information is used to progressively update and refine the surrogate model. The updated surrogate model is then employed to propose a new task condition to be presented to the subject in the next iteration by maximizing a predefined *acquisition function*. Such acquisition functions are often defined by an automatic transition from explorative to exploitative search behavior when optimization is successful (Brochu et al., 2010); this allows for an efficient and reliable search over an exhaustive task space. The aim of the study was to identify the cognitive tasks that maximally dissociated the two spatially overlapping FPNs; therefore, the “target” brain state we optimized for was the difference in evoked blood-oxygen-level dependent (BOLD) signal of the dorsal over ventral FPN (dorsal FPN > ventral FPN).

### Tower of London and Deductive Reasoning tasks maximally differentiate the networks

In Stage 2, the optimization algorithm successfully searched the 2D-task space (Supplementary Fig. 1a). Across all subjects, we observed that both the Tower of London and the Deductive Reasoning task were the most frequently sampled tasks by the acquisition function (Fig. 2a). Such sampling behavior is highly reflective of identified optima: once the algorithm’s uncertainty about the experiment space decreases as the run progresses, it transitions into an exploitative phase, in which the acquisition function keeps sampling the predicted optimum or nearby in the task space. Equally, when estimating a group-level Bayesian model based on all observations, the Tower of London and Deductive Reasoning tasks were predicted to be optimal for disambiguating the two networks (Fig. 2b). This result was highly consistent within and across subjects (Supplementary Fig. 2) and could not be explained by the particular spatial arrangement of the tasks (Supplementary Results). Post-hoc analyses, assessing spatial similarity of the group-level predictions within different sub-regions of the dFPN and vFPN, confirmed that neuroadaptive Bayesian optimization chose tasks that selectively activated the entire dFPN, and not only sub-regions (Fig. 3a).

**Figure 2:**
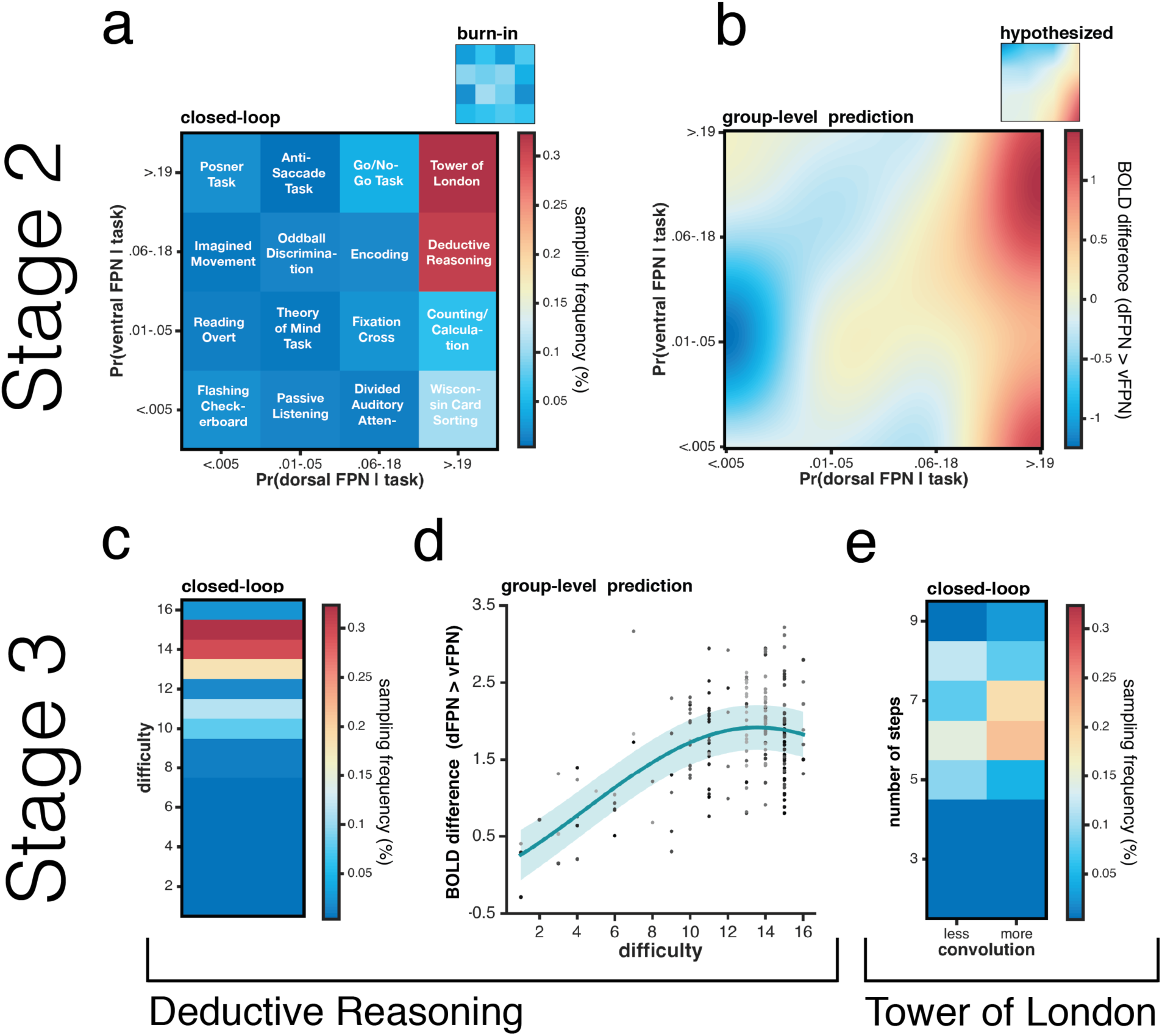
Group-level results. **(a)** Real-time sampling behavior of neuroadaptive Bayesian optimization and **(b)** group-level predictions across task space identify the Tower of London and Deductive Reasoning task to be optimal for dissociating the dFPN from the vFPN. **(c)** Real-time sampling behavior of optimization algorithm and **(d)** group-level predictions (observations colored by subject) show difficulty levels 13 to 15 to be optimal in Deductive Reasoning task. **(e)** Real-time sampling behavior of optimization algorithm for Tower of London task suggests 6 to 7 steps to be optimal.

**Figure 3:**
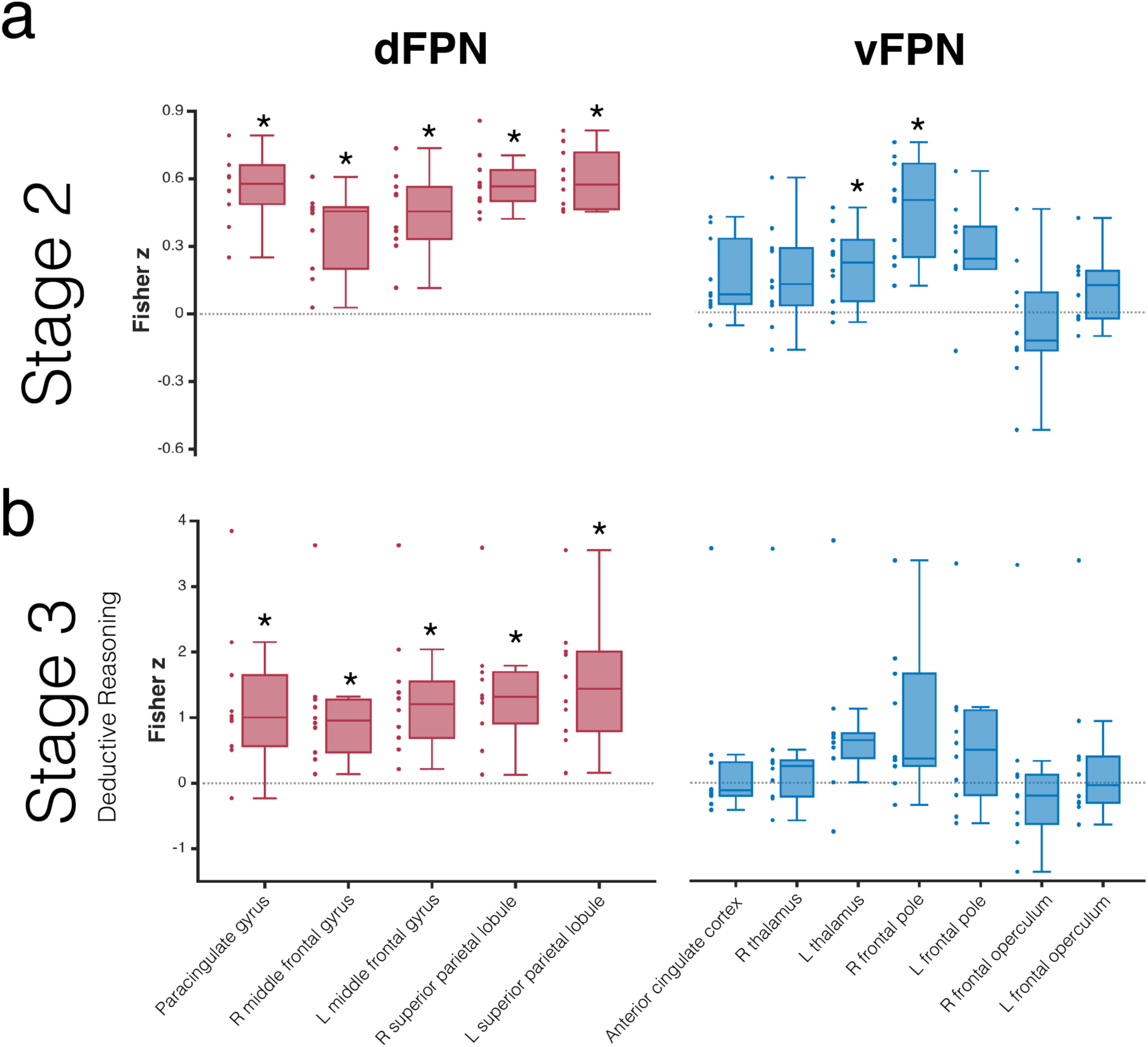
Post-hoc cluster-based analyses within dFPN and vFPN. To demonstrate that the optimization method selected tasks (for Stage 2) and task parameters (for Stage 3) that selectively targeted the entire dFPN, we performed post-hoc whole-brain back-projections (see Methods). Asterisks indicate significant findings after correcting for multiple comparisons using permutation testing (critical *t(*9) = 3.46 for Stage 2; critical *t*(9) = 2.63 for Stage 3). **(a)** We find that for Stage 2 identified tasks optimally increased the contrast with the vFPN across the entire dFPN, i.e., our obtained real-time results were not driven by single clusters within the dFPN. In comparison, most regions within the vFPN did not differ from zero; this was expected (since we contrasted beta-coefficients from each voxel’s time series with the beta-coefficients of the vFPN, see Methods). (**b)** This result was replicated for the Deductive Reasoning task in Stage 3. This analysis was not carried out for the Tower of London task because the group-level predictions were different from the subject-level results (see Supplementary Fig. 4). On each box, the central mark is the median, the edges of the box are the 25^th^ and 75^th^ percentiles, the whiskers extend to the most extreme datapoints considered not to be outliers. Left to each plot are all individual observations (*n*=10 for both Stages).

### Optimal tasks are contrary to predictions by existing meta-analysis

The finding that the Tower of London and Deductive Reasoning tasks best segregated the dorsal and ventral FPN was inconsistent to the initial meta-analysis (Yeo et al., 2014) that predicted the Wisconsin Card Sorting and Counting/Calculation tasks to be optimally suited for this purpose (Fig. 2b - small panel). This unexpected finding was formally assessed by comparing the spatial similarity between voxel-level Bayesian predictions and the, according to the meta-analysis, hypothesized predictions across the whole experiment space. This analysis confirmed that the meta-analysis was significantly different from the obtained group-level results within the whole dFPN (*t*(9)= 5.62, *p*<.001, paired two-tailed t-test). For a more focused analysis, the same analysis was performed for five separate clusters within the dFPN. In line with the previous result, this analysis revealed that for each cluster within the dFPN the obtained group-level results were significantly more likely than the hypothesized predictions (Fig. 4).

**Figure 4:**
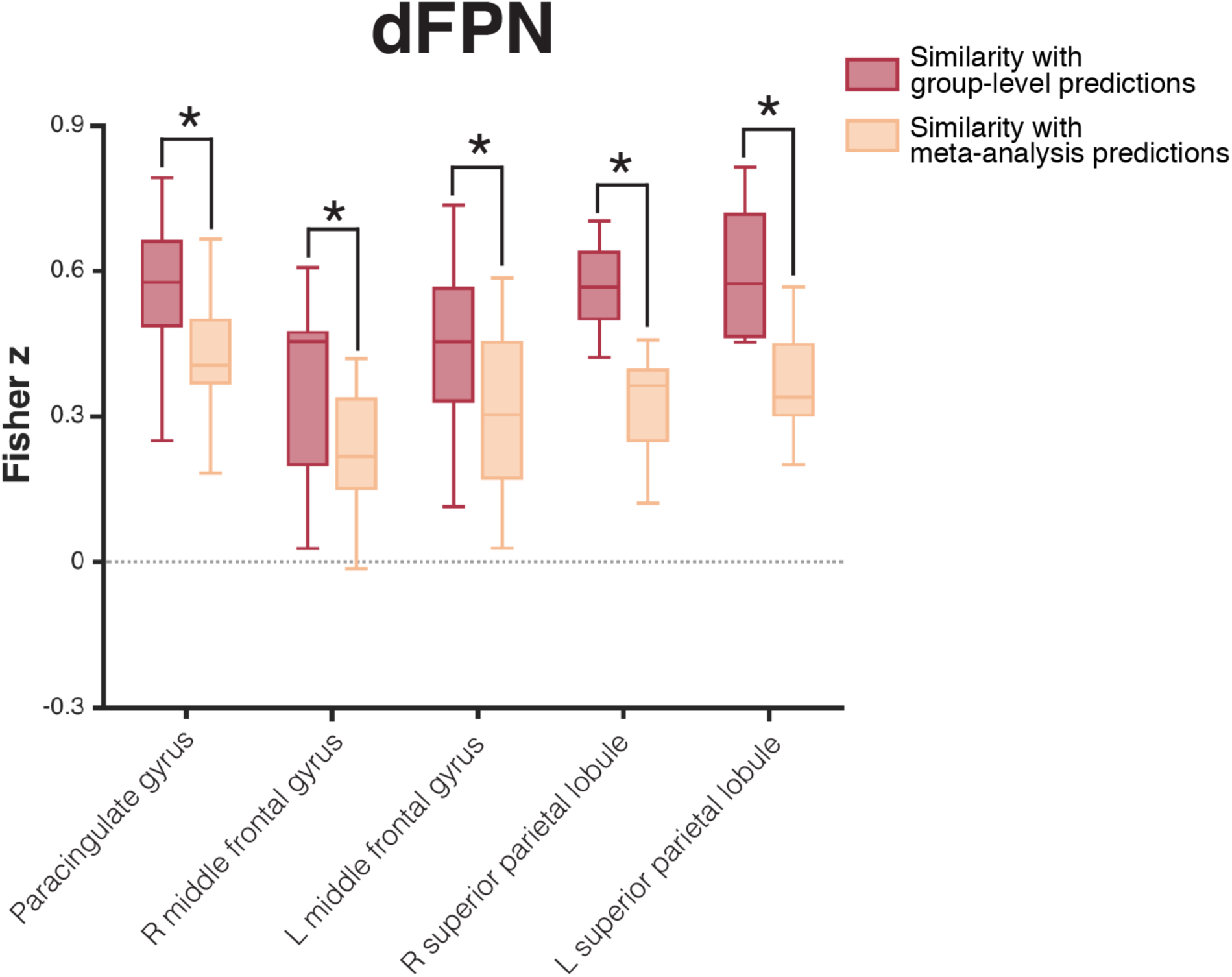
Post-hoc cluster-based comparison between group-level predictions and hypothesized predictions derived from meta-analysis. In order to statistically assess differences between our obtained group-level Bayesian predictions and the, according to the meta-analysis, hypothesized predictions across the whole task space, we compared two different post-hoc whole-brain back-projections (see Methods). Asterisks indicate significant findings after correcting for multiple comparisons using permutation testing (critical *t*(9)=2.81). We found that for each cluster within the dFPN the obtained group-level results were significantly more likely than the hypothesized predictions. On each box, the central mark is the median, the edges of the box are the 25^th^ and 75^th^ percentiles, the whiskers extend to the most extreme datapoints considered not to be outliers.

### Activity profiles across many tasks necessary to segregate large-scale brain networks

In an additional post-hoc analysis, we contrasted activation between the dFPN with the other large-scale networks (Yeo et al., 2014). This resulted in qualitatively distinct patterns of activity across the task space for the different network contrasts (Fig. 5). We observed that no single task on its own is able to differentiate activity between all networks, consistent with the existence of a many-to-many mapping between brain networks and cognitive processes (Price & Friston, 2005). However, at the same time, the profile of activity across the whole task space is more unique to the different networks, providing a more robust way to differentiate between networks.

**Figure 5:**
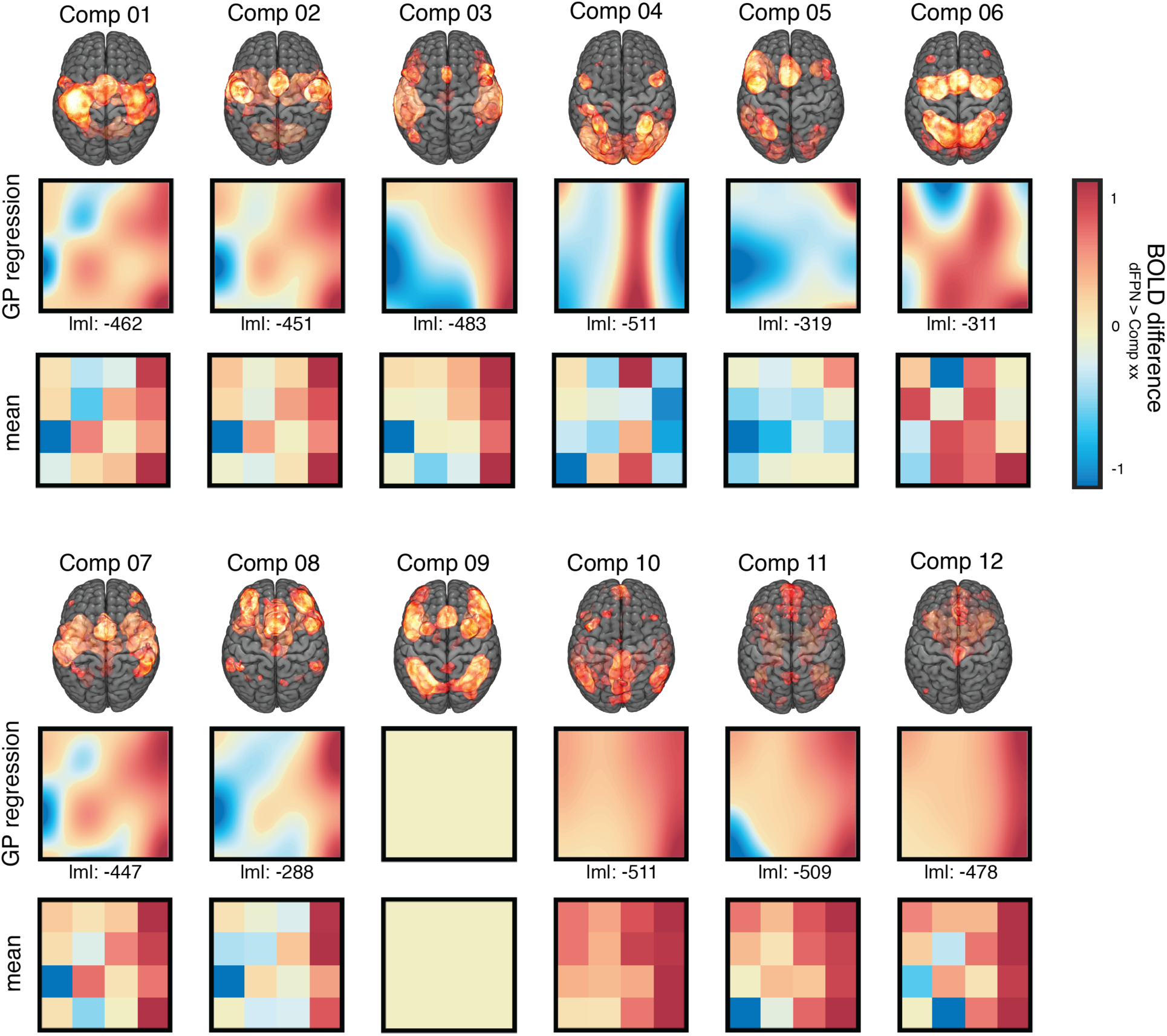
Activity patterns between dFPN and other large-scale networks. For this analysis, we contrasted brain activity of the dFPN (i.e., Comp09) with all 12 components identified in the meta-analysis. Contrast values were fed into the Bayesian optimization algorithm to derive Bayesian prediction across the whole task space for each component (‘GP regression’ panels). Log marginal likelihood (lml) values indicate model evidence of the Bayesian predictions, with higher values implying higher model evidence. To explore if the regression resulted in sensible predictions, we also plotted the mean value across all subjects for each cell of the task space (‘mean’ panels). This analysis yielded qualitatively distinct network activity patterns. We observed that no single task on its own is able to differentiate activity between all networks, consistent with the existence of a many-to-many mapping between brain networks and cognitive processes. However, at the same time, the profile of activity across the whole task space is more unique to the different networks, providing a more robust way to differentiate between networks. It should be noted that the task space was designed for the contrast of dFPN > vFPN (i.e., Comp08); this is also reflected in the highest model evidence value (lml) for this contrast; thus, other contrasts should be interpreted with caution.

Importantly, in many ways (with the exceptions noted in the previous section) our findings were consistent with the meta-analyses (Yeo et al., 2014), demonstrating how neuroadaptive Bayesian optimization can be used to discover both where meta-analyses are broadly correct as well as highlight divergence. Notably, Comp02, a network encompassing the supplementary motor area that was associated with “Overt Recitation/Repetition” and “Overt Reading” in the meta-analysis showed differential activation (i.e., Comp02 > dFPN in blue) for the Overt Reading task also in the present study. Comp3, a network including the auditory cortex, was predicted to be associated with “Pitch Monitoring/Discrimination” and “Passive Listening” in the meta-analysis and showed high activation for Passive Listening and the Divided Auditory Attention Task in this study. The left lateralized network, Comp05, including the inferior frontal gyrus (i.e., overlapping with Broca’s area) has been linked to “Covert Naming” and “Semantic Monitoring” in the meta-analysis and showed pronounced differential activation for Reading, Theory of Mind, Counting/Calculation and Encoding tasks, with the latter tasks also involving processing of verbal information, such as numbers. The finding from the meta-analysis that tasks preferably recruiting Comp06 were “Saccades” and “Anti-Saccades” was consistent with our results, in which the highest differential activation for Comp06 was also associated with the Anti-Saccade task. Furthermore, networks encompassing structures of the default mode network, such as Comp10, Comp11 and Comp12, were mainly deactivated during task performance. It should be noted that the task space was designed for the contrast of dFPN > vFPN; this is also reflected in the highest model evidence value (lml, see Methods) for this contrast (i.e., dFPN > Comp08); thus, other contrasts should be interpreted with caution. In summary, this analysis suggests that for a comprehensive understanding of how the dFPN differs from other networks (than just the vFPN), it is desirable to consider activity across the whole task space.

### Increased relational integration separates FPNs

In Stage 3, the parameters of the optimal tasks from Stage 2 were fine-tuned, producing a network dissociation that was even finer-grained. For the Deductive Reasoning task we successfully optimized (Supplementary Fig. 1b) across a 1D-space with 16×1 difficulty levels (see Methods). Across all subjects, difficulty levels 13 to 15 were most frequently selected by the acquisition function during the real-time period (Fig. 2c). This result was confirmed when estimating a group-level Bayesian model (Fig. 2d) and was highly consistent across all subjects (Supplementary Fig. 3). Linear mixed-effect (LME) analyses indicated a significant quadratic relationship between difficulty and brain activity (*χ*^2^(1) =12.07, *p*<.001, likelihood ratio test), suggestive of plateauing performance with difficulty (Fig. 2c/d and Supplementary Fig. 6). Difficulty levels 13 to 15 encoded Deductive Reasoning problems involving high relational integration, indicating that such cognitive operations may drive the optimal differentiation between the dFPN and vFPN. Assessing spatial similarity of the group-level predictions within separate sub-regions of the dFPN and vFPN, confirmed that these task parameters selectively activated the entire dFPN (Fig. 3b).

### Multi-step planning separates FPNs

The Tower of London task was successfully optimized (Supplementary Fig. 1c) across an 8 (number of steps) × 2 (convolution) parameter space (see Methods). Across all subjects, spatial planning problems involving 6 to 7 steps were most frequently selected by the acquisition function during the real-time period (Fig. 2e). When estimating a group-level Bayesian model for the Tower of London task, distinct predictive patterns for the two convolution stages across the experiment space were observed (Supplementary Fig. 4a); however these effects seemed to be driven by a few individuals (Supplementary Fig. 4b). This assumption was confirmed when conducting a LME analysis that showed a significant quadratic effect of number of steps with brain activity (*χ*^2^(2) =76.18, *p*<.001, likelihood ratio test) but no significant effect of convolution (*p*=.92) or interaction (*p*=.95). Assessing the predicted optimal task parameters on a subject-level again indicated inter-subject reliability (Supplementary Fig. 5).

## Discussion

In summary, we have discovered and refined two tasks that both strongly dissociate a dorsal from a ventral FPN. Interestingly, these results were not predicted by the previous meta-analysis (Yeo et al., 2014). From a cognitive perspective, the Deductive Reasoning task (at the higher difficulty levels) involved increasingly complex relational integration. The optimal Tower of London task involved multi-step planning. Across both tasks the dissociation between the FPNs was less sensitive for convolution or multiple, yet non-integrated rules; such integration processes have been associated with frontopolar networks in the past (Bunge, Wendelken, Badre, & Wagner, 2005; Christoff & Gabrieli, 2000; Parkin, Hellyer, Leech, & Hampshire, 2015).

In the present study, neuroadaptive Bayesian optimization allowed an efficient closed-loop search through a large task space, which would have been intractable with standard neuroimaging methodology. Importantly, we found consistent replicable results across and within each individual subject; this is not a trivial advance in light of the growing concerns about the reproducibility of fMRI findings (Button et al., 2013; “Fostering reproducible fMRI research,” 2017; Ioannidis, Munafò, Fusar-Poli, Nosek, & David, 2014; Munafò et al., 2017; Szucs & Ioannidis, 2017). Moreover, the technique transitioned from exploration to a focused search, thereby including an inbuilt replication stage, as it repeatedly kept sampling the predicted optimum by collecting new data (i.e., data previously unseen by prior analysis). This is in line with the recent advocacy of “out-of-sample” prediction in experimental sciences (Bzdok & Yeo, 2016; Yarkoni & Westfall, 2016).

In contrast to meta-analyses, neuroadaptive Bayesian optimization is less biased by prevailing theories of the field, requires far less data and can explore effects that meta-analyses are blind to, such as specific contrasts of interest, functional connectivity or unsampled regions in the task space. Here, we demonstrated that the meta-analysis provides a useful starting point, and that it summarizes many of the network activity profiles well; however, when disambiguating dorsal from ventral FPNs, the meta-analysis provides a partially misleading picture. The discrepancy between the meta-analysis and our results could have arisen for a number of reasons. For example, it could because of the absence of direct comparisons among different tasks in meta-analyses, differences in baseline or contrast conditions. This needs to be studied in further detail in future work; however, the current approach highlights the importance of studying a large number of tasks within the same analysis and testing framework.

The post-hoc analyses showed that in order to achieve a comprehensive understanding of how the dFPN differs from other networks (than just the vFPN), it is necessary to consider activity across the whole task space. This implies that when simultaneously disambiguating between many networks requires batteries of cognitive tasks rather than a single task. This is important not just from a theoretical point of view, but also when designing cognitive assessment batteries that are aimed at sensitively discriminating network function, e.g., in developmental or clinical settings.

The findings that seemingly different cognitive processes (planning and relational integration) are chosen to differentiate dorsal and ventral FPNs may indicate that the neurobiological processes do not align well with the pre-existing cognitive taxonomy. One possibility is that the neuroadaptive Bayesian optimization has discovered a neurobiological discrete process common to both planning and reasoning that is not normally discriminated in assessments. An alternative explanation is that the two tasks share some additional sub-component unrelated to planning and reasoning, that is driving the difference in activity between the networks. Either way, the results highlight the importance of combining insights from neurobiology and behavior when developing a cognitive taxonomy.

In conclusion, the results of the present study have demonstrated the powerful synergy between neuroadaptive Bayesian optimization and meta-analyses for research question within the cognitive neurosciences that historically have been challenging to tackle. It was shown how neuroadaptive Bayesian optimization can go beyond the meta-analysis: using it as a starting point for exploration, before optimizing tasks at a far finer precision than otherwise possible.

## Methods

### Subjects

21 healthy volunteers (13 female, 28.67 ± 5.11 years, range: 21-41 years, 3 left-handed) took part in the real-time optimization study; of these, 10 participated in Stage 2, and 11 in Stage 3. For Stage 3, one subject was excluded due to excessive head movement for both runs. Mean frame-wise displacement (FD) (Power, Barnes, Snyder, Schlaggar, & Petersen, 2012) of this subject in both runs was 0.22 mm and 0.23 mm, respectively; both runs were identified as significant outliers using an iterative implementation of the Grubbs test (*α* = 0.05) (Barnett & Lewis, 1994). Subjects gave written informed consent for their participation. The study was approved by the Hammersmith Hospital (London, UK) research ethics committee. All subjects had no history of any neurological/psychiatric disorders and had normal or corrected-to-normal vision. Subjects were informed about the real-time nature of the fMRI experiment but no information was given on the actual aim of the study or which parameters would be adapted in real-time. The investigator was not blinded due to the complexity of data acquisition and the need to ensure that real-time optimization was functioning.

### FPN dissociation

In Stage 1, the target brain state was defined based on the meta-analysis reported in Yeo et al. (Yeo et al., 2014) In this study, 12 cognitive components were identified based on 10,449 experimental contrasts covering 83 BrainMap-defined task categories. The authors made publicly available the brain maps that contain the probability of components activating different brain voxels, i.e. *Pr*(voxel | component), as well as the probability for each task recruiting those components, i.e. *Pr*(component | task). We focused on two components within the multiple demand cortex (Duncan & Owen, 2000): Component 8, a ventral frontoparietal network (vFPN) and Component 9, a dorsal frontoparietal network (dFPN). Thresholded (z > 1) and binarized maps of the two components were used as networks of interest for both studies. The target brain state we optimized for in both studies was the difference in brain activity between these two networks (i.e., dFPN > vFPN).

### Experimental procedure

In Stage 2, subjects underwent two separate real-time optimization runs. Task space and target brain state were identical across the two runs; both runs were initiated randomly (i.e., the first five blocks were selected randomly from across the task space). For one subject, we only conducted one run due to a failure in the projection to the MR screen. Tasks were presented in a block-wise fashion for 35 s followed by 19 s of rest (black background). Preceding each task, participants received a brief instruction (5 s) about the task they would need to perform in the upcoming block followed by a short rest period (3 s). Each run automatically ended after 20 blocks (20.67 min). For two subjects, one out of the two runs stopped after only 13 blocks due to technical failure.

In Stage 3, subjects again underwent two separate real-time optimization runs; this time optimizing the tasks that were predicted to be optimal in Stage 2. The target brain state was identical to Stage 2 (i.e., dFPN > vFPN). One run optimized for the Deductive Reasoning task while the other run optimized for the Tower of London task (see tasks description below). The order of runs was counter-balanced across participants. Each run automatically ended after 20 blocks (18 min). Before each run started, we informed the subjects via microphone which of the two tasks they would need to perform in upcoming run.

Subjects were trained and familiarized with all tasks outside of the scanner prior to the start of the experiment. All tasks were implemented in Matlab using Psychophysics Toolbox (Brainard, 1997; Pelli, 1997). Auditory stimuli were presented using sound-attenuating in-ear MR-compatible headphones (Sensimetrics, Model S14, Malden, USA).

#### Deductive Reasoning Task

The Deductive Reasoning task in Stage 3 was based on an implementation by Hampshire et al. (Hampshire et al., 2012). A 3×3 grid of cells was displayed on the screen and each cell contained an object. Each object was made up of three different features: color, shape and number of copied. The features were related to each other according to a set of rules. The subject had to deduce the rules that relate the object features and identify the object whose contents did not correspond to those rules. Task difficulty increased along a single dimension (16×1) with increased complexity of the rules applied. Deductive Reasoning problems were presented for 30 s followed by a 5 s response interval, in which subjects were presented with two objects (i.e., the correct one and a randomly selected one), between which subjects chose by pressing the right or left button on a button response box.

#### Tower of London Task

Equally, the Tower of London task in Stage 3 was based on an implementation by Hampshire et al. (Hampshire et al., 2012). Numbered beads were positioned on a tree shaped frame. The participant were instructed to reposition the beads mentally so that they were configured in ascending numerical order running from left to right and top to bottom of the tree. Task difficulty increased along two dimensions (8×2); the first dimension represented the total number of moves required (from 2 to 9 moves); the second dimension represented the convolution operation (more or less). Convolution was defined as, when for the participant to succeed on the task a correctly placed bead must first be displaced. Tower of London task problems were presented for 30 s followed by a 5 s response interval, in which subjects were presented with two numbers (i.e., the correct number of moves and the correct plus or minus one number of moves), between which subjects chose by pressing the right or left button on a button response box.

### Real-time fMRI

Images with whole-brain coverage were acquired in real-time by a Siemens Verio 3 T scanner using an EPI sequence (T2*-weighted gradient echo, voxel size: 3.00 × 3.00 × 3.00 mm, field of view: 192 × 192 × 105 mm, flip angle: 80°, repetition time (TR)/echo time (TE): 2000/30 ms, 35 interleaved slices). Prior to the online run, a high-resolution gradient-echo T1-weighted structural anatomical volume (voxel size: 1.00 × 1.00 × 1.00 mm, flip angle: 9°, TR/TE: 2300/2.98 ms, 160 ascending slices, inversion time: 900 ms) and one EPI volume were acquired. Online (as well as subsequent offline) pre-processing were carried out with FSL (Jenkinson, Beckmann, Behrens, Woolrich, & Smith, 2012). The first steps occurred offline prior to the real-time fMRI scan. Those comprised brain extraction (Smith, 2002) of the structural image followed by a rigid-body registration of the functional to the downsampled structural image (2 mm) using boundary-based registration (Greve & Fischl, 2009) and subsequent affine registration to standard brain atlas (MNI) (Jenkinson, Bannister, Brady, & Smith, 2002; Jenkinson & Smith, 2001). The resulting transformation matrix was used to register the two networks of interest from MNI to the functional space of the respective subject. For online runs, incoming EPI images were motion corrected (Jenkinson et al., 2002) in real-time with the previously obtained functional image acting as reference. In addition, images were spatially smoothed using a 5 mm FWHM Gaussian kernel. For each TR, means of the dFPN and vFPN masks were extracted. The second stage of pre-processing involved removing large signal spikes with a modified Kalman filter (Koush, Zvyagintsev, Dyck, Mathiak, & Mathiak, 2012). The pre-processed timecourses were written into a separate text file for subsequent analyses. Removal of low-frequency linear drift and movement artifacts was achieved by adding a linear trend predictor and six motion parameters as confound regressors into the general linear model (GLM). The experiment commenced after 10 TRs to allow for T1 equilibration effects.

### Neuroadaptive Bayesian Optimization

#### Target measure

After presentation of each task block, we calculated the difference in brain level activation between the two target brain networks. For this purpose, we ran incremental GLMs on the pre-processed timecourses of each target network separately. In our case, incremental GLM refers to the design matrix growing with each new task block, i.e., the number of timepoints as well as the number of regressors increasing with the progression of the real-time experiment. The GLM consisted of task regressors of interest (one regressor for each block), task regressors of no interest (e.g., 5 s instruction period in Stage 2 or 5 s response interval in Stage 3) as well as confound regressors as described above (six motion parameters and linear trend) and an intercept term. Task regressors were modeled by convolving a boxcar kernel with a canonical double-gamma hemodynamic response function (HRF). After each new block, the beta coefficients were re-estimated. We then computed the difference between the estimates of all task regressors of interest for the two networks (dFPN > vFPN) and entered them into the Bayesian optimization algorithm. An initial burn-in phase of five randomly selected experimental conditions was employed in both studies, i.e., the first GLM was run after the first five blocks after which the closed-loop experiment commenced.

#### Bayesian model

In the first step of Bayesian optimization, all available samples acquired up to that iteration are used to update the algorithm’s surrogate model by predicting the brain’s response across the entire experiment space (also for unseen points) using Gaussian process (GP) regression (Brochu et al., 2010; Rasmussen & Williams, 2006; Shahriari et al., 2016). GP-based surrogate models are commonly used in Bayesian optimization due to their flexibility and tractability (Brochu et al., 2010; Shahriari et al., 2016). GPs are fully specified by their mean and covariance functions. As a prior, we employed a zero mean function and as covariance function we chose the squared exponential (SE) kernel (Rasmussen & Williams, 2006). The SE kernel encodes the basic prior assumption that points close in the task space elicit similar brain responses while points far from each other could exhibit distinct responses:

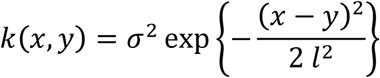

where *x*, *y* ∈ ℝ^2^ correspond to the choice of experimental condition. The hyper-parameters σ ∈ ℝ and *l* ∈ ℝ^2^ each determine the variance and length scale of the covariance kernel, respectively. Further, it is assumed that observations are corrupted by white noise, 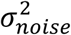. These parameters need to be selected prior to running the experiments. We used independent data from four/two subjects for Stage 2/Stage 3 to tune these parameters using Type-2 maximum likelihood (Rasmussen & Williams, 2006). This choice of hyper-parameters was then fixed for all subjects’ runs.

#### Acquisition function

In the second step of Bayesian optimization, by maximizing a pre-defined acquisition function, a new point within the experiment space is selected to sample from in the next iteration. In our study, we used the expected improvement (EI) acquisition function that automatically trades-off between exploration and exploitation (Brochu et al., 2010) by studying the predictions of the surrogate model as well as the uncertainty of the predictions. Informally, this choice of acquisition can be seen as trying to maximize the expected improvement (i.e., the target measure) over the current best. We define *m*(*x*) as the predictive mean for a point *x* ∈ ℝ^2^ and *std*(*x*) as the predictive standard deviation. The expected improvement is defined as (Brochu et al., 2010):

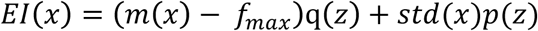

where *q*() and *p*() are defined as the cumulative and probability density functions for a standard normal distribution respectively and f_max_ can be either the maximum predicted or maximum observed value of the objective function depending on our assumptions about the noisiness of the observations (Brochu et al., 2010). Finally, z is defined as:

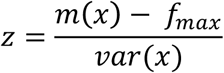

At every iteration, the next experimental condition to be observed is selected by maximizing the expected improvement:

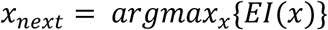

In Stage 2, f_max_ was set to be the maximum predicted value. As in Stage 3 we observed much more exploitative behaviour of the acquisiton function for the first three subjects; therefore, we set f_max_ for the final seven subjects to be the maxmimum observed value which lead to increased exploration.

### Real-time sampling behavior of optimization algorithm

The EI acquisition function is characterized by an automatic transition from explorative to exploitative search behavior when optimization is successful. This results in the acquisition function exploring the experiment space in the beginning of the run, when uncertainty is the greatest, by subsequently proposing experimental conditions far away in the experiment space. Once the algorithm’s uncertainty about the experiment space decreases as the run progresses, it transitions into an exploitative phase, in which the acquisition function keeps sampling the predicted optimal experimental condition or nearby in the experiment space. In order to assess if the optimization successfully transitioned from exploration to exploitation in our study, we computed the mean ± std Euclidean distance (ED) between successive experimental conditions across all subjects over course of the run; this distance should decrease with the transition from exploration to exploitation, suggesting successful optimization. Results of these analyses are depicted in Supplementary Fig. 1. In addition, we computed the sampling frequency of each experimental condition for the closed-loop period of the experiment (i.e. iteration 5-20); this contrasts with the burn-in period (i.e., iteration 1-5) in the beginning of the experiment when five experimental conditions were selected randomly. The results of these analyses are depicted in Fig. 2 and provide insight about the experimental conditions that were predicted to be optimal across the whole group.

### Bayesian predictions across the experiment space

In order to obtain group-level Bayesian predictions across the whole experiment space, we conducted GP regression based on all available observations of all subject. Hyper-parameters of the group-level GPs were automatically retuned using Type-2 maximum likelihood (Rasmussen & Williams, 2006). The result of the analyses are depicted in Fig. 2b/d and Supplementary Fig. 4 and reveals the experimental conditions that are predicted to be optimal across the whole group.

To better account for subject-specific effects in Stage 3, we conducted linear mixed effect (LME) models. Data entering the LME models were subject-level Bayesian predictions across the task parameters (Supplementary Fig. 3/5). For each of the 10 subjects this yielded in 16×1 Bayesian predictions for the Deductive Reasoning and 8×2 Bayesian predictions for the Tower of London task. We constructed LMEs either with a linear or quadratic term for linking Bayesian predictions of brain activity with difficulty/number of steps for the Deductive Reasoning/Tower of London task. For the Tower of London task, we added two additional terms: one linking Bayesian predictions of brain activity with convolution and one interaction term (number of steps × convolution). We then conducted likelihood ratio tests to compare the goodness of fit between the two models (linear vs. quadratic model). Results of this analysis are reported in the Results section.

### Post-hoc cluster-based analyses within dFPN and vFPN

To demonstrate that the optimization algorithm had successfully selected tasks that targeted the entire dFPN, we conducted post-hoc whole-brain back-projections. For this, we compared the spatial similarity of the group-level Bayesian predictions across the task space (Fig. 2b-large panel) with Bayesian predictions calculated from the timecourse of each voxel in the brain. This analysis was done on offline-preprocessed data in each subject’s native space. Data underwent motion correction (Jenkinson et al., 2002), spatial smoothing (5 mm FWHM), linear de-trending, and scrubbing (Power et al., 2012) (FD threshold > 1.5) before GLMs were conducted on each individual voxel time series. Resulting beta-coefficients from each voxel’s time series were then contrasted with beta-coefficients derived from the vFPN timecourse (as we were interested in the voxel-wise dissociations from the vFPN); and these contrast values were fed into the GP regression (the Bayesian optimization’s model) to derive Bayesian predictions across the whole task space for each voxel. Hyper-parameters of the GP for each voxel were automatically selected using Type-2 maximum likelihood (Rasmussen & Williams, 2006). The Bayesian predictions for each voxel were then spatially correlated with the group-level Bayesian predictions, resulting in one similarity value (Pearson *r*) per voxel. Whole-brain similarity maps were subsequently Fisher z-transformed and registered to MNI standard space. For Stage 2, within-subject runs were averaged before further analyses were conducted. Next, dFPN and vFPN maps were thresholded at *z* > 2 in order to obtain separate clusters within each network. This resulted in five clusters for the dFPN (i.e., paracingulate gyrus, right middle frontal gyrus, left middle frontal gyrus, right superior parietal lobule and left superior parietal lobule) and seven clusters for the vFPN (i.e., anterior cingulate cortex, right thalamus, left thalamus, right frontal pole, left frontal pole, right frontal operculum, and left frontal operculum); clusters with less than 200 voxels were not considered. For each cluster mask, mean z-values of each subject (*n*=10) were subsequently extracted. Group-level inference was carried out on that data using a one-sample two-tailed t-test, and permutation testing was used to correct for multiple comparisons (using t_max_-method (Blair & Karniski, 1993); number of possible permutations was 2^10^). Results of this analysis are depicted in Fig. 3.

### Post-hoc cluster-based comparison with meta-analysis

We performed statistical inference to compare group-level Bayesian predictions (Fig. 2b-large panel) with the meta-analysis-based hypothesized predictions (Fig. 2b-small panel) for the entire task space. The same processing pipeline was used as described above; however the Bayesian predictions of each voxel were correlated with the hypothesized meta-analytic predictions, again resulting in one similarity value (Pearson *r*) per voxel. These whole-brain similarity maps were subsequently Fisher z-transformed and registered to MNI space. For the whole-network analysis, we extracted mean z-values within the dFPN mask for both similarity maps (i.e., hypothesized predictions and group-level predictions) in each subject, and performed a paired two-tailed t-test (results reported in Results section). For a more focused analysis, we extracted mean z-values for all five clusters within the dFPN separately in each subject (same clusters as described above), and performed a paired two-tailed t-test on these values, with permutation testing to correct for multiple comparisons (using t_max_-method (Blair & Karniski, 1993); number of possible permutations was 2^10^). Results of this analysis are depicted in Fig. 4.

### Post-hoc activity patterns between dFPN and other large-scale networks

An additional post-hoc analysis assessed the differential activity patterns between the dFPN and other large-scale brain networks across the whole task space. For this analysis, timecourses of thresholded (*z* > 1) and binarized maps from all 12 components identified in Yeo et al. (Yeo et al., 2014) were extracted from the offline-preprocessed data. GLMs were conducted on each component’s timecourse and resulting beta-coefficients were contrasted with beta-coefficients from the dFPN (i.e., component 9). These contrast values were fed into the GP regression to derive Bayesian prediction across the whole task space for each component (Fig. 5 – *GP regression*). Hyper-parameters of the GP for each component were automatically selected using Type-2 maximum likelihood (Rasmussen & Williams, 2006). We also computed log marginal likelihood (lml) values, indicating model evidence of the Bayesian predictions (with higher values implying higher model evidence). To validate, if the GP regression resulted in sensible predictions, we also plotted the mean value across all subjects for each cell of the task space (Fig. 5 - *mean*). For visualization purposes, for each component, we re-scaled all positive contrast values between 0.1 and 1 and all negative contrast values between -1 and 0. This procedure had the benefit of keeping important information about the “sign” of the contrast values while facilitating plotting using an identical scaling throughout all subplots.

### Code availability

For GP regression, we use a Python implementation from: http://github.com/SheffieldML/GPy. Python code for different acquisition functions is available from: http://github.com/romylorenz/AcquisitionFunction.

## Acknowledgement

This work was supported by the NIHR Imperial BRC and the Leverhulme Trust. I.R.V. is funded by the Wellcome Trust (103045/Z/13/Z).

